# Computational Urban Ecology of New York City Rats

**DOI:** 10.1101/2025.07.21.665423

**Authors:** Ralph E. Peterson, Dmitry Batenkov, Ahmed El Hady, Emily L. Mackevicius

## Abstract

Urban rats are highly adaptable, thriving in the dynamic and often inhospitable conditions of modern cities. Despite substantial mitigation efforts, they remain an enduring presence in urban environments, yet surprisingly little is known about the daily lives and behavioral strategies that underlie their success. Here, we conducted fieldwork on free-ranging rats in New York City, using thermal imaging and ultrasonic audio recordings. We apply cutting-edge artificial intelligence techniques to capture high-resolution movement patterns and generate 3D reconstructions of foraging environments including subways, streets, and parks. We characterize social vocalizations across environmental contexts, and compare the patterns of social communication observed in NYC rats to the distribution of rodent vocalizations reported in the literature. This work provides a foundation for translating techniques and theories of rodent cognition from the lab to urban ecological settings.

## 1 Introduction

The study of animal cognition in complex urban environments is increasingly tractable, and increasingly timely, due to advances in techniques and theories for understanding cognitive behavior. Studying animals, especially rodents, in urban environments has a strong history in ethology and ecology [9, 17, 25, 38, 49, 57, 72], and new methods are emerging in neuroscience [33]. An understanding of behavior in urban contexts is crucial for mitigation efforts, designing cities, and controlling disease spread. A recent revolution in the development of artificial intelligence approaches to understand behavior (e.g. keypoint tracking and action recognition) have enabled analysis of animal kinematics at unprecedented spatiotemporal resolution, and these approaches are beginning to be deployed in the field [11, 37, 45, 51, 77]. Moreover, machine learning-based techniques to quantify acoustics (e.g. animal vocalizations, soundscapes) are coming online [18, 61, 63, 65]. Despite progress in these domains, quantification of animal behavior in the wild remains technically challenging. For example, tracking multiple animals in the lab is non-trivial in part due to animals occluding each other’s body parts during social interactions; this problem is further complicated *en natura* by occlusions from the environment (e.g. leaves, branches, rocks, etc.). In the acoustic domain, the ability to segment, featurize, and classify environmental sounds is a persistent problem under active research. This is primarily due to the complexity and variability of real-world soundscapes, which often contain overlapping sources, non-stationary noise, and lack large-scale annotated datasets [64]. As neuroscientists uncover the neural basis of cognitive behaviors in lab settings, there is increasing interest in studying the neural basis of an expressive set of natural behaviors [13, 16, 21, 32, 40, 41, 47, 55]. We must develop integrated behavioral analysis systems and multimodal modeling frameworks that are robust in the environmental complexities of the real world. Here, we address this gap by performing fieldwork in wild New York City (NYC) rats and introducing a computational toolkit for analysis of behavior and the environment.

Rats have been reported in NYC since early colonial days, and have experienced intense selective pressure since then. Rats originally became commensal with human cities in Asia a couple thousand years ago. After reaching Europe around 800 years ago, brown rats (*Rattus norvegicus* — also called “Norway rats” due to a misconception about their origins) rapidly became a prominent urban pest. They spread throughout Africa, the Americas, and Australia during the 18th and 19th centuries as a result of European colonialism [42]. Rat populations likely experienced strong selective pressure within port cities, which underwent rapid industrialization and urbanization [22]. A recent study [20] used isotopic analysis and mass-spectrometry to analyze archaeological (1550s–1900 CE) rat remains from eastern North America and provided a large-scale framework for species arrival, inter-specific competition, and dietary ecology. The study found that brown rats arrived earlier than previously expected and rapidly out-competed black rats in coastal urban areas. Urban environments changed dramatically from the late 19th into the 20th century, a period that spans around 500 rat generations. An analysis of the genomes of New York City brown rats found intriguing signatures of adaptation near genes associated with metabolism, diet, the nervous system, and locomotor behavior ([22]), and there is evidence for a significant change in rat cranial shape in New York City over a 120-year period [56].

Rats demonstrate a remarkable ability to adapt to the changing urban environment. Rats only require a single ounce of food and water a day to live [6]. They primarily find food and shelter at human habitations, and therefore they must interact with humans in various ways. In particular, the city’s rats adapt to practices and habits among New Yorkers for disposing of food waste [70]. Curbside overnight garbage disposal from residences, stores, subway and restaurants, as well as littering, contribute to the sustenance of the city’s rats. Rats nearly always use the same routes to their food sources, probably using olfactory cues or their own secretions such as urine [69]. Interestingly, rat behavior changed during the COVID lockdown, as their access to food was altered [4, 50].

Urban rats face many of the same challenges as human city dwellers. One striking commonality between urban humans and rats is their diet [19]. Today, the urban human diet contains an increasingly large proportion of highly processed sugars and fats that cause a number of public health concerns. Some of these health concerns could conceivably apply to rats as well. Rats in New York City play a very important role in spreading disease, and are commonly infested with fleas, lice and mites that carry bacteria that can cause disease in humans, including bubonic plague, typhus and spotted fever [15]. Bartonella pathogens (which can cause cat scratch disease, trench fever, Carron disease) and various viruses were also found in New York City’s rats in this study. Our study of the behavioral repertoire, movement patterns and collective behaviors of New York rats could add an important layer of understanding of how diseases spread via the behavioral dynamics of rats, informing mitigation efforts.

Understanding rat behavior, and its relation with particular features of urban environments, is a crucial piece of rat mitigation efforts. Various municipalities have declared a “war on rats” since at least the early 1900s, and current best practices involve integrated pest management measures, which mitigate environmental conditions that attract and support rats, instead of relying heavily on rodenticide poison [35]. The NYC Health Department offers “Rat Academy” training programs that teach pest control professionals about rat behavior and habitats, and how environmental features contribute to rat infestations [44]. Our AI-driven approach could contribute to these control measures by developing a computational understanding of how rat behavior is shaped by environmental conditions, and rapidly identifying areas of interest for pest control in a dynamic manner.

From the perspective of understanding cognition, laboratory rats are one of the most important animal models for both basic and translational research [75]. A key motivation of this study is to unravel the heterogeneous behavioral repertoires exhibited by wild rats in urban environments such as New York City. We transfer advanced analytical and AI-driven methods, developed to study animals in the lab, to the study of rats in the wild. Using these methods, we characterize the foraging behavior of NYC rats at a high spatiotemporal resolution, focusing on coordinated rapid movement of groups of rats, 3D geometry of foraging environments, and vocal communication.

## 2 Results

### 2.1 Targeting field recordings of rats in NYC

Rats are abundant across NYC (Figure 1A). Manhattan, the borough with the highest density of reported rat sightings, has approximately 200 reports of rat per square mile per year (Figure 1B), and across NYC, the NYC311 complaints center receives approximately 50 reported rat sightings per day (Figure 1C). There is a seasonality to rat sightings, with more rats reported in summer months (Figure 1D). Thus, we collected data throughout July 2024 from three distinct locations in Manhattan. We used public data [43, 73] to identify specific candidate sites, and set out to record the foraging behavior of groups of NYC rats using RGB and thermal imaging cameras, as well as ultrasonic microphones (Figure 2A). Rats prefer to forage in the evening, and in shadowed areas, which poses a challenge for standard videography methods, but thermal videography made it possible to visually resolve rodents, even in dark shadowed areas (Figure 2B). Using these methods, it was possible to observe groups of rats foraging in a variety of urban environments, including subways (Figure 2C), city streets (Figure 2D), and parks (Figure 2B), including even areas with some occlusion due to fencing and underbrush (Figure 2E).

**Figure 1:**
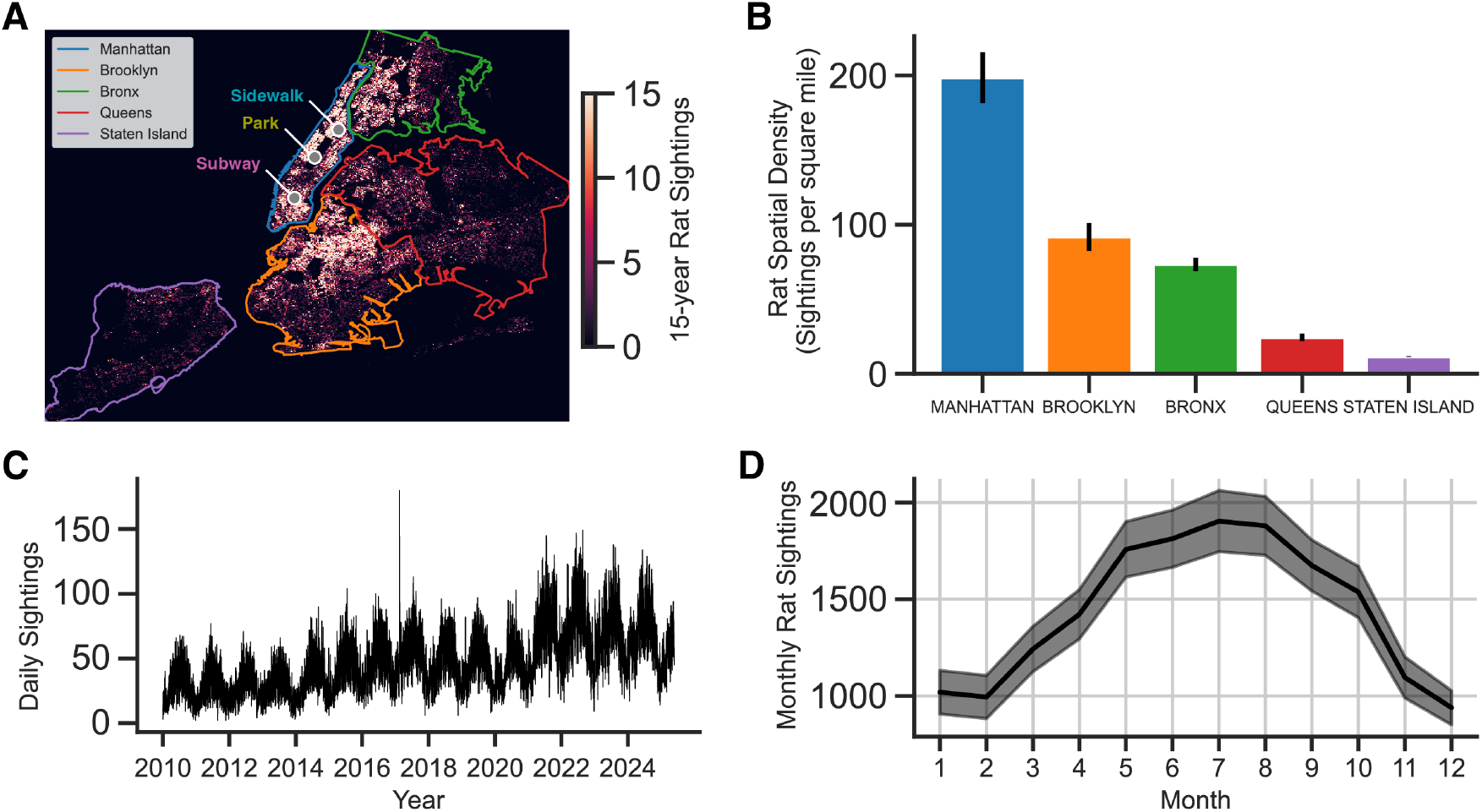
New York City citizen reports of rat sightings. **(A)** Spatial heatmap of rat sightings over a 15-year period generated from NYC311 reports. Recording locations from the study are labeled with gray points (Subway, Park, Sidewalk). **(B)** Spatial density of rat sightings computed by dividing the total number of rat sightings (yearly) by the land area of each borough. Error bars are SEM, computed across the 15-year period. **(C)** Daily rat sightings over 15-year period shows cyclic rat activity. **(D)** Number of sightings by month shows seasonality of sightings, with reports peaking in Summer months. Error bars are SEM, computed across the 15-year period.

**Figure 2:**
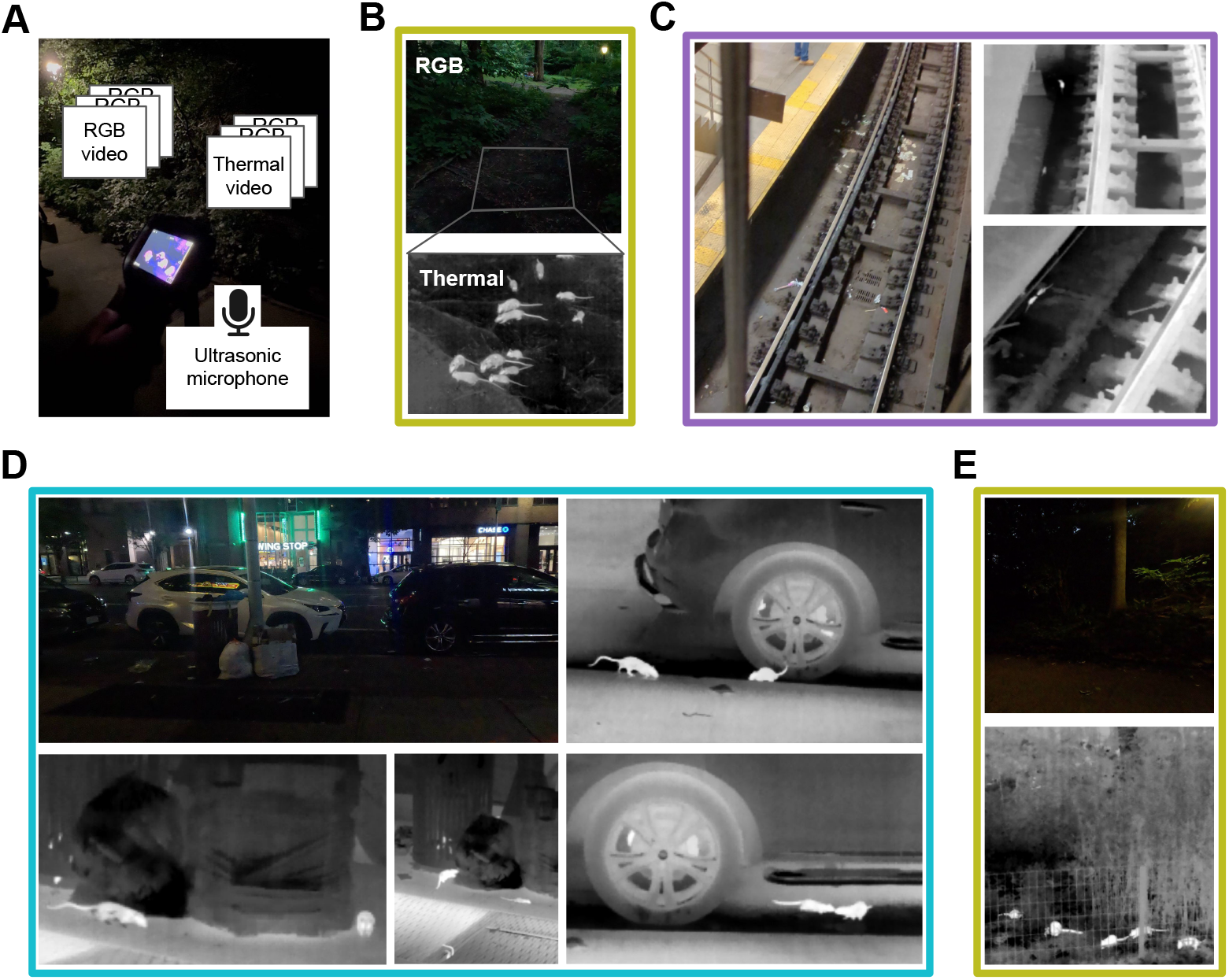
Groups of rats foraging in NYC environments observed using thermal imaging. Colored outlines correspond to sidewalk, park, and subway sites indicated in Figure 1A. **(A)** Photo of data acquisition setup, with overlayed schematic of data sources **(B)** (top) RGB image of an area where rats were observed foraging in Central Park, New York, NY. Gray outline indicates field of view for thermal imaging camera. (bottom) Frame from a thermal video of rats foraging in the area outlined above. **(C)** RGB image of a subway scene in NYC where rats were observed foraging, together with selected thermal video frames showing rats. **(D)** RGB image of a street scene in NYC where rats were observed foraging, together with selected thermal video frames showing rats. **(E)** RGB image of a park scene at night with dense underbrush behind a fence, and thermal image of rats foraging in this scene.

### 2.2 High-resolution tracking of real-world rat movements using thermal imaging

While multi-object tracking is a well-researched area in AI and computer vision [36], and in particular in the context of animal tracking [34, 37, 52, 60, 71, 77], applying the different existing approaches in our field study to extract accurate rat tracks from thermal videos (Figure 3A) proved to be a nontrivial task. In outdoor real-world environments, specific challenges include varying number of animals per frame, a wide distribution of animal sizes, multiple animal types (e.g. rats and squirrels), occlusions, and inaccurate and missing detections. In particular, extracting object boundaries (masks or bounding boxes) rather than skeletons was desirable for size estimation. Finally, to process large amounts of videos, we were interested in methods providing near-realtime (or faster) processing speeds. Our detection and tracking pipeline (Figure 2B) included the most recent version of the YOLO (You Only Look Once) model [30, 59], fine-tuned on 50 hand-labeled frames from our thermal videos of rats foraging, combined with the ByteTrack tracking algorithm [78], which is robust to occlusions and missing track segments, and utilizes low-score detections and Kalman filtering for predicting new locations.

**Figure 3:**
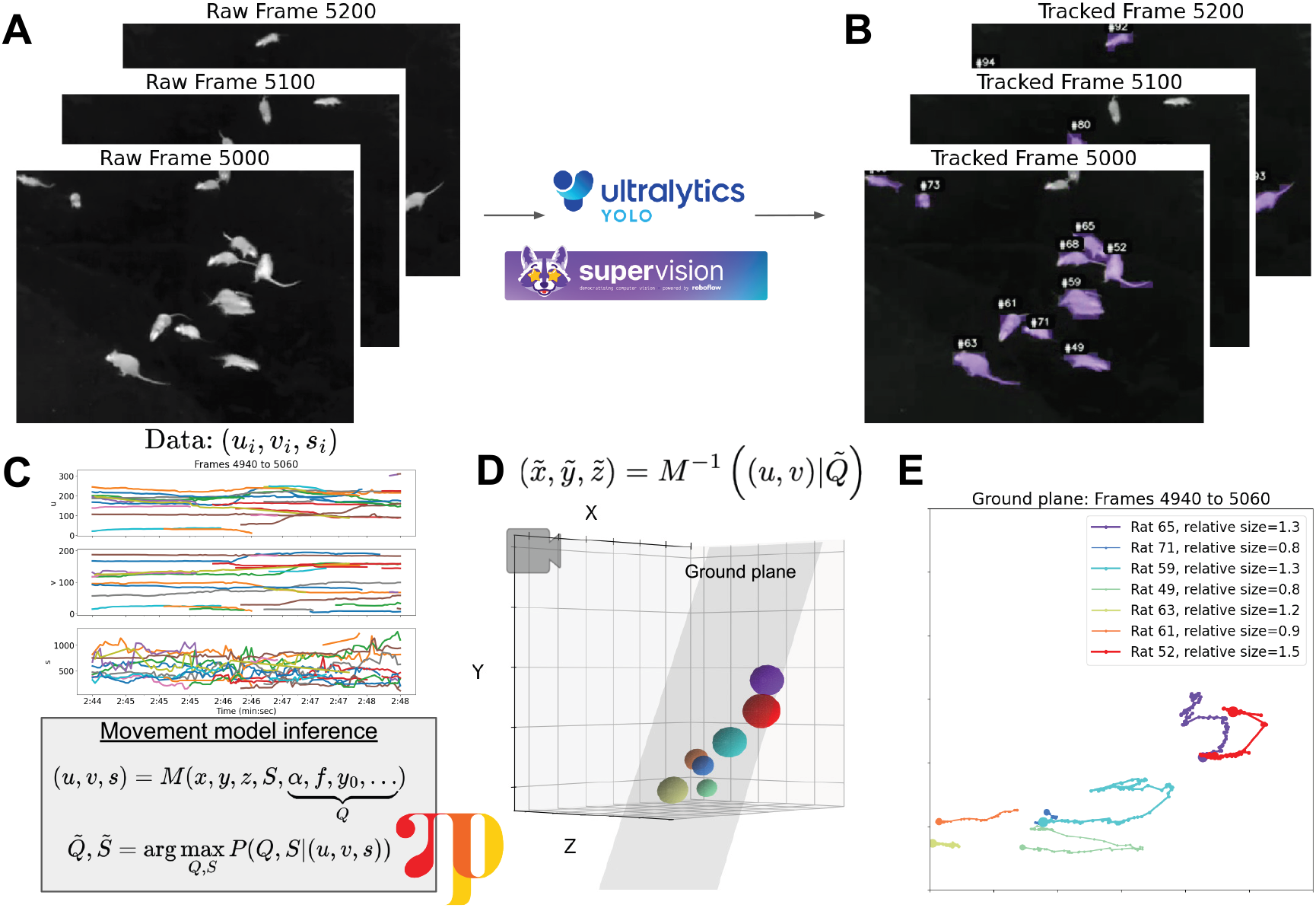
Tracking rats and inferring their relative size and 3D position. **(A)** Raw thermal video frames showing rats of a range of sizes **(B)** Thermal video frames overlayed with bounding masks of rats detected with YOLO **(C)** (top) Tracked position (*u, v*) and size *s* of rats in pixel-space (bottom) Model relating true 3D position and relative size (*x, y, z, S*) to position and size in pixel-space, further described in Section 4.2 **(D)** Inferred 3D positions and relative sizes of rats, shown in 3D for a single frame, and **(E)** projected onto the ground plane for a series of frames.

Interestingly, some rats, likely juveniles, appeared to be substantially smaller than others. The pixel position (*u, v*) and bounding box size (*s*) of each rat was computed over time (Figure 3C, top). However, the video was recorded at an angle tilted relative to the ground plane. As a result, mask sizes in pixel space are disproportionately larger when rats are closer to the camera. We wondered whether we could infer the true relative sizes and 3D positions of rats from the raw track data. We constructed a simple probabilistic model *M* relating, for each tracked rat *i*, its true 3D position 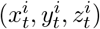 at frame *t* and its true size (*S*^*i*^), to camera-pixel position 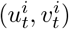 and apparent size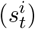. This model depends on intrinsic and extrinsic camera parameters (*Q*), under the assumption that rats move on a ground plane facing the camera at some angle, and approximates each rat by an on-plane circle. Using the probabilistic programming language Pyro [5], 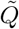 and 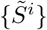 were inferred from the observed data (Figure 3C, bottom). These inferred parameters were subsequently used to estimate rats’ ground plane positions throughout the video (Figure 3D,E) by a straightforward projection. Relative size is computed in units of the bounding box area, and normalized by dividing by the average rat’s size.

In order to compare the movements of rats of different sizes, we analyze ‘tracklets’, where each tracklet corresponds to a trajectory of a rat from when it is first detected until it is lost (e.g. it exits the frame). Tracklets shorter than 5 seconds are excluded. We analyze a 10 minute video of rats foraging, with 201 extracted tracklets and the corresponding relative sizes estimated by the procedure described above. Note that each rat may enter and exit the frame multiple times, so there are more tracklets than true rats in the scene. In the analyzed video, the distribution of inferred animal sizes includes small rats, large rats, and a larger squirrel (Figure 4A). Larger animals appeared to move faster than smaller animals (Figure 4B-C). Groups of rats sometimes appeared to move in coordination. We quantified these moments of coordinated group activity by computing how many rats were moving faster than a minimum speed threshold (Figure 4D). Interestingly, rats that participated in moments of coordinated movement tended to be larger than non-participants (Figure 4E).

**Figure 4:**
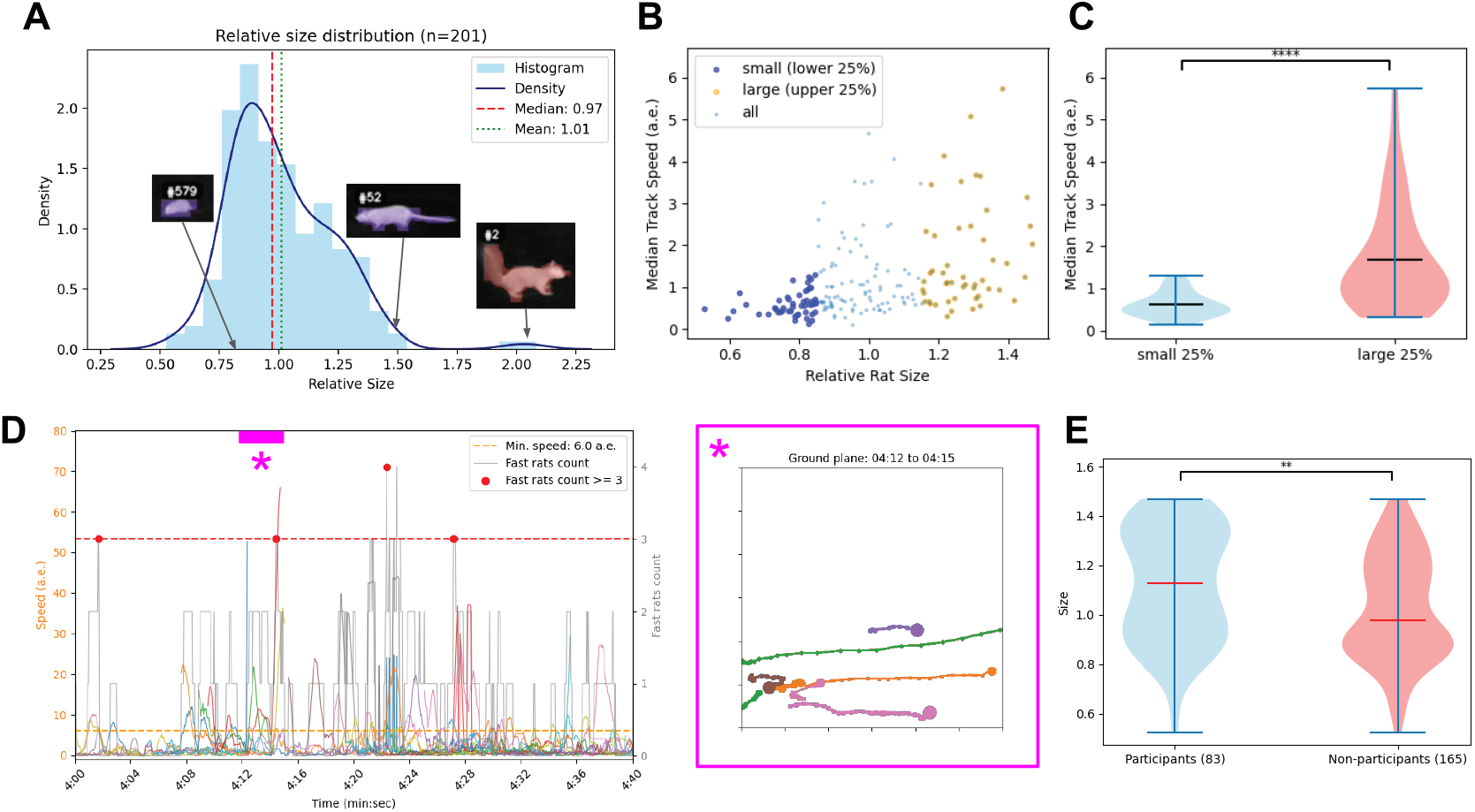
Comparing movements of smaller and larger rats. **(A)** Distribution of inferred relative sizes for all tracklets. Insets show examples of a small rat, a large rat, and a squirrel. **(B)** Median speed of each tracklet as a function of inferred relative size **(C)** Distributions of speed for tracklets in the lower and upper size quartiles (two-sample Kolmogorov–Smirnov statistic 0.56, p-value 1.4 × 10^*−*7^).**(D)** Speed as a function of time for a segment of the video, showing individual rats’ trajectories, as well as the number of fast rats. Detected moments of coordinated movement are indicated by red dots. Magenta inset shows inferred trajectories of rats during one particular epoch. **(E)** Distributions of rat sizes for rats that participate in coordinated movements, and non-participants (two-sample Kolmogorov–Smirnov statistic 0.222, p-value 0.007)

### 2.3 Capturing environmental geometry and statistics using gaussian splatting

We wondered whether it was possible to quantitatively capture 3D models of environments in which rats were foraging. Recent advances in computer vision make it possible to reconstruct 3D models of a scene from a sparse collection of camera angles [29, 39]. Urban environments where rodents forage pose some potential challenges, because they can be highly dynamic spaces, with shadowed areas, and limited access. We wondered whether it was possible to reconstruct 3D models of such environments using data acquired with a single handheld RGB camera. Multiple camera views were collected of a park scene (Figure 5A), and run through the gaussian splatting model, which first involves estimating camera positions using the COLMAP algorithm (Figure 5B), then constructing a 3D model in the form of collection of 3D gaussians that best explain the data. These models capture environmental geometry at a high level of detail (Figure 5C). The models also allow for quantification of aspects of the environmental statistics that may be relevant to rodent foraging, such as the degree of shelter versus open-field. We found that this could be captured by analyzing the standard deviation of gaussian centers in the z-axis (Figure 5D). Gaussian splats were generated for a diverse range of urban foraging environments, ranging from parks to streets to subways. Figure 5E-H shows the same analysis pipeline as in Figure 5A-D, for a subway scene, using images captured from an overpass above the subway platform.

**Figure 5:**
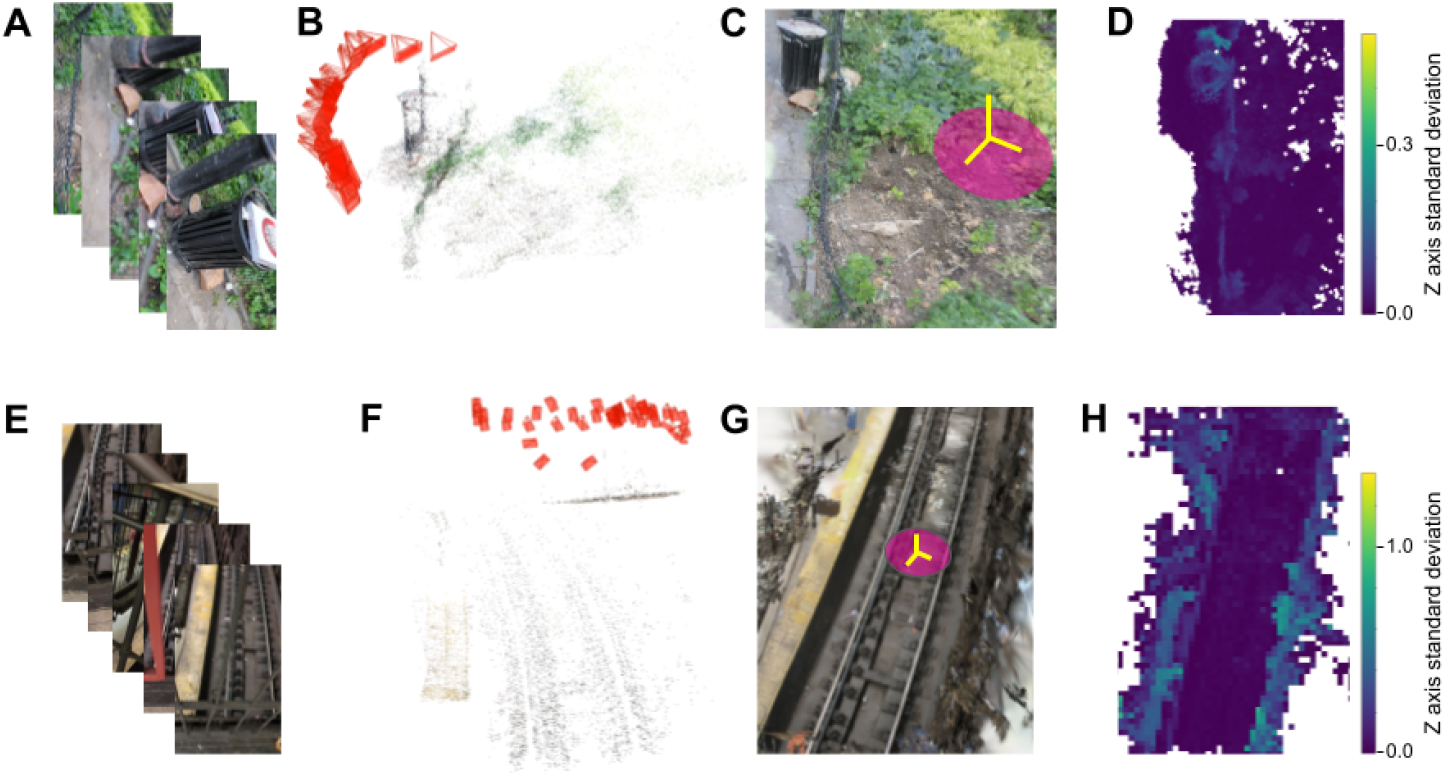
Capturing environmental geometry and statistics using gaussian splatting. **(A)** Example RGB images of a park scene from different angles. **(B)** Inferred camera positions and points in the 3D scene, from the COLMAP algorithm. **(C)** 3D scene composed of gaussians inferred by the gaussian splat algorithm. **(D)** Top-down heatmap of this 3D scene, showing the standard deviation in the Z axis of gaussian locations. **(E-H)** Same as **(A-D)**, for a subway scene.

### 2.4 Ultrasonic acoustic recordings of rat vocalizations in New York City

There have been few attempts to document the social vocal lives of rats in their natural urban habitat. Here, we used a wireless ultrasonic microphone (Figure 6A) to record vocalizations emitted by rats in different urban environments during social interactions. First, we find that environments substantially differ in their acoustics (Figure 6B). For example, the Subway environment is approximately 12 dB (4x amplitude) louder than the Park environment. Next, we extracted vocalization annotations from raw audio using a deep neural network (Deep Audio Segmenter, [68]). Vocalizations mostly occurred in bouts (sequences of vocalizations separated by less than 250 ms of silence) and were observed in various types of social interactions (Figure 6C-H).

**Figure 6:**
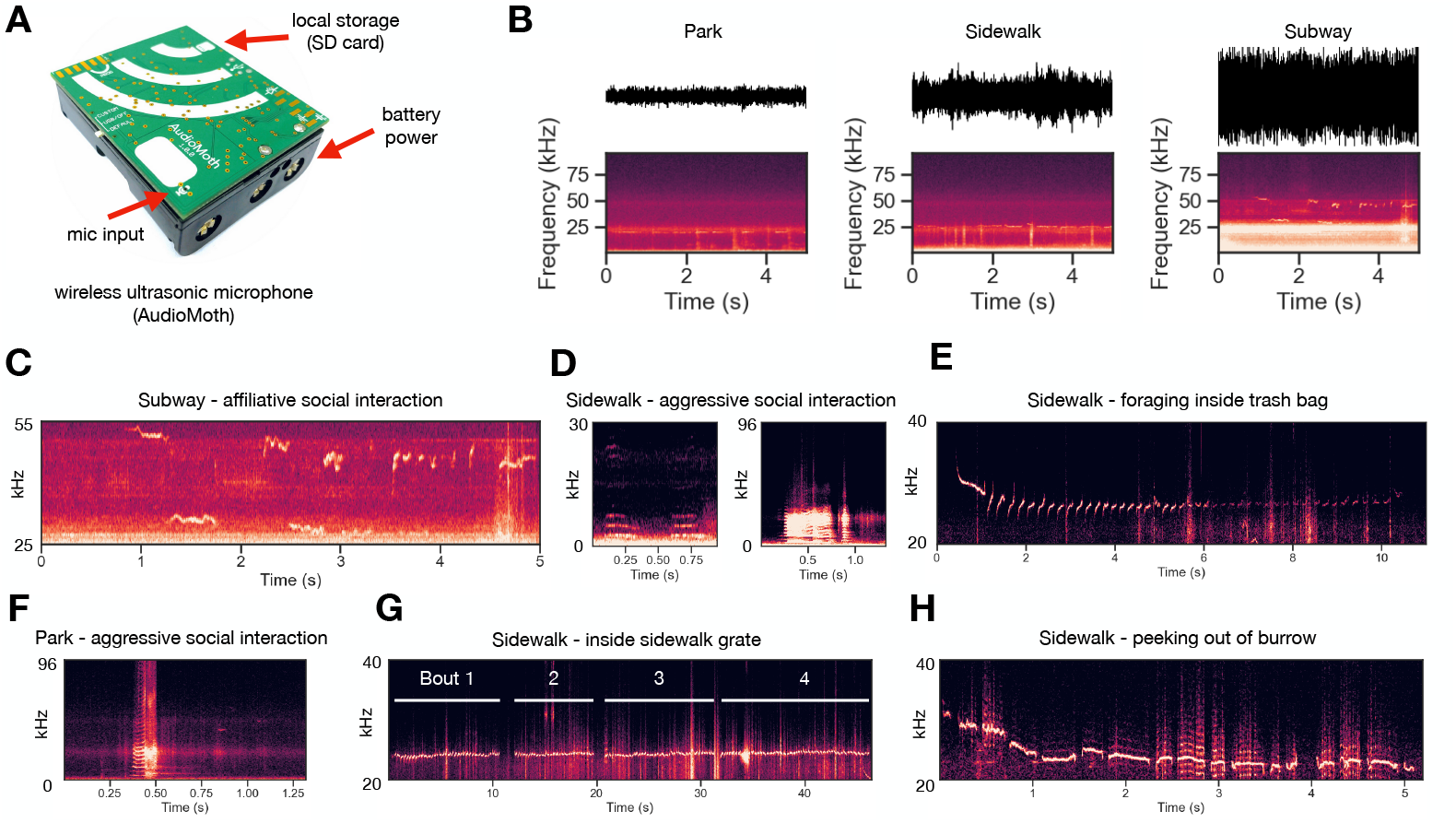
Ultrasonic acoustic recordings of NYC rat vocalizations in diverse soundscapes. **(A)** Field recordings were taken using an Audiomoth wireless ultrasonic microphone. **(B)** Recordings were taken from three distinct environments (Subway, Park, Sidewalk) which vary acoustically. **(C-H)** Vocalization spectrograms from distinct social contexts and environments. **(C)** Affiliative social interaction between two rats in the subway tracks. **(D)** Human-audible vocalizations from an aggressive interaction between pairs of rats on the sidewalk nearby trash cans **(E)** Long-duration bout of calls recorded from a rat foraging inside of a trashbag on the sidewalk. **(F)** Human-audible vocalizations from an aggressive interaction between a pair of rats in the park. Note classically affiliative USVs between 0.75-1.25s. **(G)** Multiple bouts of vocalizations recorded from a sidewalk grate where rats were frequently seen entering and exiting. **(H)** Rat cautiously poking its head out of a burrow hole from a sidewalk tree lawn.

Next, we calculated acoustic features of vocalizations and compared them to a large-scale meta-analysis of previously published rat vocalization features. Figure 7A shows an example vocalization bout (same as Figure 6E) with DAS-annotated onsets and offsets of every syllable in the bout (purple). Next, we estimated the fundamental frequency at each time point in the spectrogram (Figure 7B) and calculated the median for downstream analysis. We computed the median frequency +/- 1 standard deviation for all bouts detailed in Figure 6, then projected their measurements into a duration vs. frequency feature space detailed by [76] (Figure 7C). We find that wild NYC rat vocalizations are consistently shorter duration and lie outside of the historical frequency-duration range reported in the meta-analysis.

**Figure 7:**
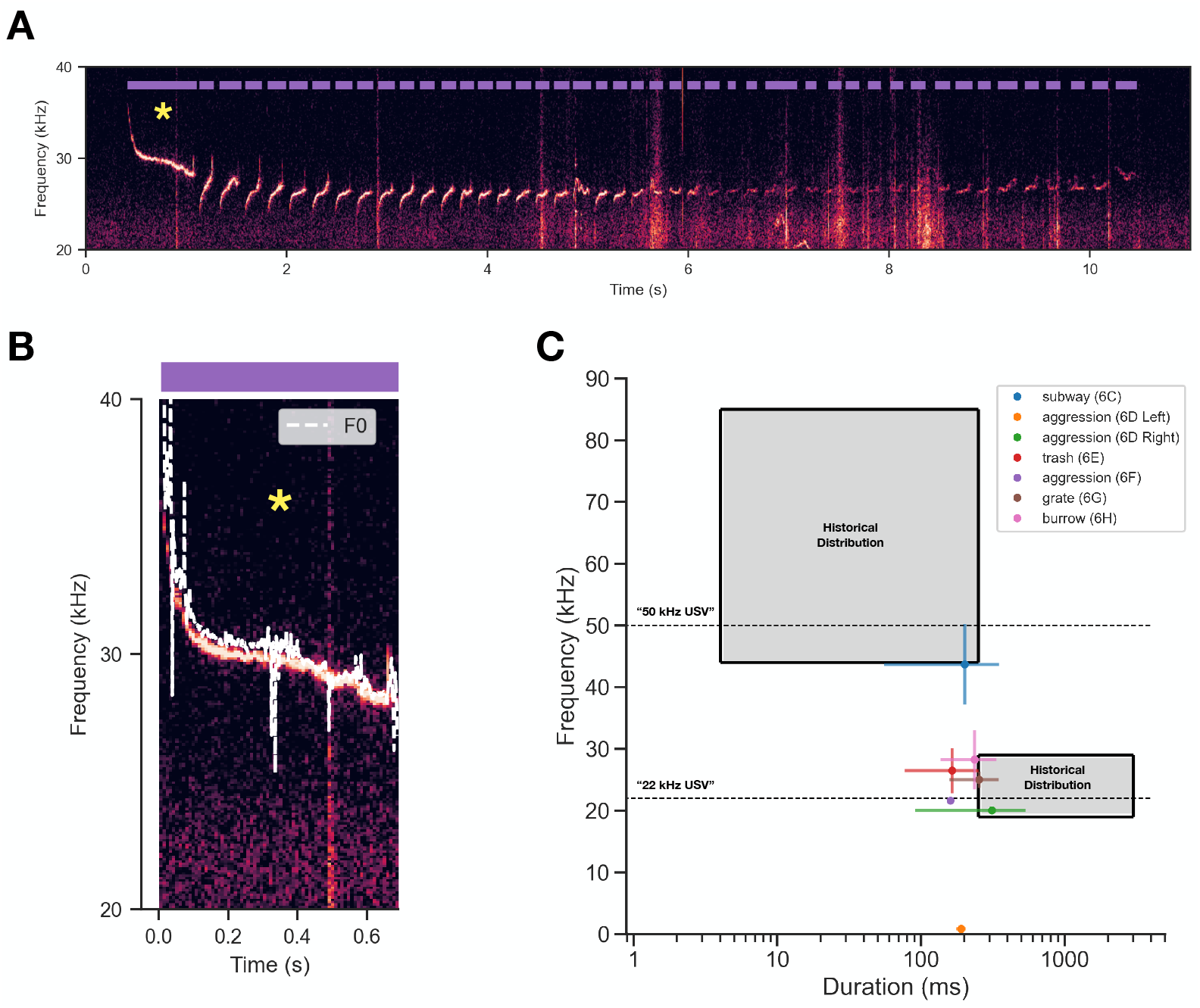
Vocalization frequency and duration are unique in NYC rats. **(A)** Same spectrogram as Figure 6E showing onsets/offsets of vocalizations extracted using Deep Audio Segmenter. **(B)** First syllable of the panel A, showing the time-varying fundamental frequency of the vocalization (F0, white dashed line). **(C)** Projection of the vocalizations described in Figure 6 into duration-frequency space detailed in [76]. Gray shaded regions denote the historical distribution of durations and frequencies reported in an expansive literature search from [76]. Horizontal dashed lines denote the two predominant vocalization types studied in rats.

## 3 Discussion

There is an increasing interest in studying animal cognition in natural habitats, especially in light of new AI tools for quantifying behavior. The advent of new tools may make it possible to translate mechanistic biological insights from laboratory based studies to natural habitats where animals live. This study validates a computational toolkit to quantify high-resolution movement and acoustic behavior of rats living in New York City. In this unconstrained urban environment, we have demonstrated a set of computational tools that allow us to track large groups of rats, estimate the relative size of individual rats, reconstruct 3D environments in which rats operate, and record ultrasonic vocalizations. The computational techniques are all open-source, and the recording technology is non-invasive with widely available hardware, which will make reproducing this study in other animals quite straightforward. We were able to map variations in foraging speeds and coordination of movements for rats of different sizes, compare 3D environmental statistics across different places where rats forage, and classify ultrasonic vocalizations rats use in different contexts, finding vocal structure distinct from the distribution of rat vocalizations commonly studied in lab environments.

Rats demonstrate an impressive ability to survive in rapidly changing urban environments, but the question of what cognitive strategies they use remains open. With sufficient data across a range of environmental conditions, it may be possible to infer cognitive strategies from unconstrained rodent foraging behavior, using a variety of recent computational techniques[1, 2, 10, 74]. It will also be important to individually tag animals to be able to follow individuals and distinct groups over extended spatio-temporal scales.

The ultrasonic vocalizations (USVs) reported here raise questions about their function in rat social behavior and challenge traditional assumptions about vocal behaviors observed in the laboratory. USVs are primarily studied in lab environments and there have been few attempts to understand vocal repertoire diversity and social function in wild rats, though the artist Brian House recorded rat ultrasonic vocalizations in NYC for a 2022 sound installation [26]. The biological function of USVs in rodents generally is still largely unknown, but since ultrasound decays rapidly with distance, one can speculate that they may be emitted in close proximity to enhance social interactions. They might also be used as a cue to localize nearby conspecifics or to report your current location like in bats [31]. An intriguing theory proposes that since ultrasound causes agglomeration of particles in the air like odorants, rodents have evolved an active sampling mechanism which couples USVs with increases in sniffing [67] to enhance the perception of pheromones. If this were true, it would help explain why we see USVs in such diverse contexts, such as within trash bags.

It is generally thought that 22 and 50 kHz vocalizations signal aversive and appetitive contexts, respectively [8]. Here, we observe that 22 kHz vocalizations are used in diverse contexts, some of which are seemingly not aversive. For example, a long bout of near-22 kHz USVs was emitted while a single rat foraged inside of a trash bag (Figure 6E). Rats have not been reported to emit 22 kHz vocalizations while foraging in laboratory settings; instead, studies have shown that 22 kHz calls actually suppress feeding behavior [7]. An alternative theory postulates that 22kHz calls could serve a security function — that is, to signal potential threat (though unperceived) in unpredictable environments [8]. Our “Sidewalk” data (Figure 6E,G,H) provide intriguing validation of this theory, where 22 kHz vocalizations are observed in highly unpredictable urban sidewalks in the absence of overt predatory threat. The acoustics of wild rat vocalizations appear to be relatively out of distribution as compared to classically recorded lab rat vocalizations, with 22 kHz vocalizations notably shorter in duration (6E,G,H; Figure 7C). Moreover, we observed numerous vocalizations that had power in the human audible range (“squeaks”) (6D,F). Future studies should focus on the social function of squeaks in wild rats, as most rat vocalization research has focused on USVs. This direction has been promising in wild mice, with recent work revealing that squeaks are genetically heritable and used seasonally [27, 28]. More recordings in more contexts over longer timescales are required to make concrete claims about the biological function of rat vocalizations in an urban setting.

Our study calls for a better mapping of physical and soundscape environments, to better understand how environmental statistics impact collective behavior. We characterized three types of environments where rats forage in NYC – subways, streets, and parks. Using advanced computer vision techniques, we characterized high-resolution geometric features of the environment that may be relevant to rodents, specifically, the relative positioning of sheltered areas versus open areas. The high-resolution 3D environment models can in principle also be used to reconstruct rats-eye-view images to validate cognitive strategies in simulation, and semantically segment 3D scenes into behaviorally relevant areas [14, 58]. We also observed highly varied background soundscapes in different types of environments. Future work should focus on semantic segmentation of sound events in the soundscape as well as determining the approximate locations of sound sources in the environment [53, 61]. A crucial follow up to our study is a multimodal mapping of the sensory world from the perspective of the animal using a combination of computational and advanced recordings techniques, to investigate how animals interact with the world through their sensory filters.

NYC offers an especially compelling field study location, because of the wealth of public data and interest from the local population. NYC is a vibrant urban environment where it is possible to set up Citizen Science projects leveraging calls from 311 (the New York hot-line where you can report sighting of rats), social media posts depicting rats, or submissions of audiovisual rat data to a central database such as NYC Open Data [12] for analysis. The current study analyzes only a small amount of data, especially compared to what could potentially be acquired in NYC. There is a need for longitudinal passive monitoring [66], e.g. designated research sites, to better understand natural behavioral patterns by acquiring a large enough data set to infer what cognitive strategies rats deploy, and predict how groups of rats will behave across different conditions. There have been a few noteworthy attempts to perform fieldwork in NYC

The computational approach to urban ecology speaks to a myriad of issues facing us today, related to how dynamic environments impact our collective behavior. These questions are especially pressing, since the ways we interact in groups, in particular in cities, are changing rapidly compared to human evolutionary history [3]. Climate change adds another layer to the nonstationary nature of urban environments. Given these changes, it’s increasingly important to understand how dynamic urban environments impact human and animal populations, at the level of individual cognition, as well as group behavior. The toolkit outlined here quantifies urban environments and animal behavior at a high resolution, laying the groundwork for reasoning about how anticipated or potential changes will impact collective cognitive behavior.

## 4 Methods

### 4.1 Video recordings of groups of rats

Video recordings of groups of rats were obtained in public locations within New York City, including streets, subway stations, and parks. Due to the minimal equipment involved, recordings on streets and subway stations were consistent with generally allowed public usage of these spaces that do not require a permit. Recordings in city parks were performed under a research permit to ELM from the City of New York Parks & Recreation Natural Resources Group. This research is purely observational and does not alter or influence the biology, behavior or ecology of the study animals or other species, so does not require an IACUC (Institutional Animal Care and Use Committee) protocol, as determined by the Columbia University IACUC. Videos of rat foraging behavior were obtained using tripod-mounted or handheld FLIR E54 thermal cameras, and simultaneous RGB video recordings (from GoPro cameras). Each video recording lasted up to 45 minutes, and camera equipment was attended at all times. In addition, multi-viewpoint videos of each scene from different angles were acquired using a Google Pixel 7A camera, for subsequent gaussian splatting [29] analysis. Gaussian splat .ply files were generated from using the Polycam online tool [54].

### 4.2 Video analysis

The locations of animals within the thermal videos were tracked using the `Supervision` and `Ultralytics YOLO` [59, 62] libraries. For detection, we trained a YOLO v11 “nano-segmentation” model (2,834,958 parameters) on 40 hand-labeled frames from our example video, and validated on another 10 hand-labeled frames, attaining mean average precision of 98.1% at IoU of 0.5. The training was done on Apple MacBook M3 Pro machine, and took approximately 7 minutes (100 training epochs). The tracking was done using the ByteTrack algorithm, with the following parameter settings: track activation threshold of 0.2, lost track buffer of 90 frames (3 seconds), minimum matching threshold of 0.8 and 5 minimum consecutive frames (the original video is at 30 fps).

To infer rat sizes from observed tracks, we have implemented a simple planar movement model approximately consistent with the observed scene. A pinhole camera with focal length *f* is located at the origin, pointing in the (−1, 0, 0) direction. We suppose that the rats are moving in a ground plane, tilted at angle *α* radians relative to the camera viewing direction, with equation *Y* = −*X* tan *α* − *Y*_0_. The camera plane coordinates (*u, v*) of a spatial point (*x, y, z*) with *x* ≠ 0 are then given by

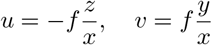

and the ground plane coordinates are given by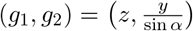. Approximating every rat by a sphere with radius *r*, the projection to the camera plane can be readily computed by the standard single view geometry transformations for quadrics (e.g. [23, Ch .8.3]) as some explicit function s of *Q* = (*α, y*_0_, *f*) and *x, y, z, r*. We take the area of this projection to be the apparent size *s*. This gives the transformation for each frame *t* and rat *i*:

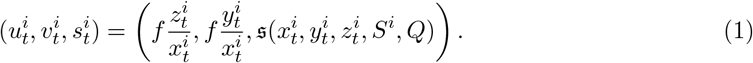

We model the movement of each rat *i* as an approximate random walk constrained to the ground plane:

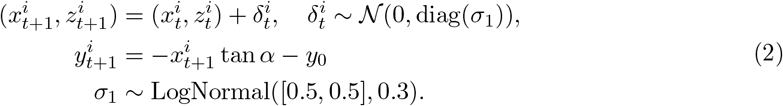

In practice, measured values of *u, v* and especially *s* are noisy, due to rats varying pose and non-spherical shapes (see Figure 3C, top panel). Therefore we also introduce latent observational noise:

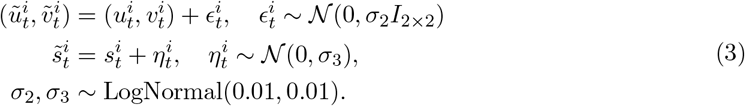

Combining (3) with (1),(2), adding simple Gaussian priors on *Q*, the initial positions 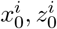 and sizes *S*^*i*^, we thus obtain a complete specification of the generative model *M*. Implementing the above model in Pyro probabilistic programming language [5], we obtain the model likelihood of the observed data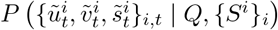. The parameters *Q*, {*S*^*i*^} are subsequently found by Maximum A-Posteriori (MAP) estimation, using Pyro’s powerful inference engine based on stochastic variational inference. This provides estimates of the rat sizes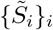. Subsequently, the 3D and the ground plane coordinates for each *i, t* are computed via

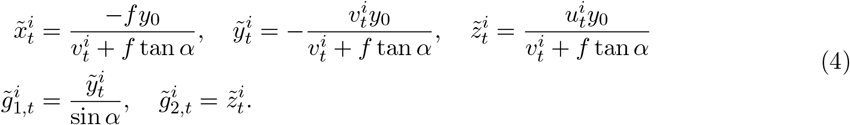

### 4.3 Audio recordings

Audio recordings were performed using a battery-powered ultrasonic acoustic logger (AudioMoth [24]). Recordings were sampled at 192 kHz in “default” mode and written as .WAV files to local storage (128 GB SD card). The files were later transferred to a workstation for analysis (see Vocalization extraction and processing). A session was acquired if rat activity was detected or suspected to be observed in a given area. The device was either handheld or placed on a surface pointed towards the area of interest at a distance of approximately 2 meters. Session durations varied between seconds to tens-of-minutes.

### 4.4 Vocalization extraction and processing

Onsets and offsets of rat vocalizations were determined using a combination of human annotation and Deep Audio Segmenter (DAS), a supervised deep-learning technique for vocalization extraction [68]. First, vocalization onsets, offsets, and type were hand annotated (n=211 USVs) and used as training data for DAS. In brief, DAS learns to predict the onsets/offsets of vocalizations in unseen audio data and performs best when given diverse training samples (e.g. vocalizations occurring in various noise conditions). Given that noise conditions are highly variable and unconstrained in urban environments, it is challenging to train a supervised model to extract vocalizations in all possible acoustic conditions. Therefore, DAS was used as an aid to generate vocalization annotations at scale, then subsequently reviewed by a human annotator. Fundamental frequency was calculated using the VocalPy python package, specifically the `vocalpy.feature.sat()` function [46]. In one instance — Figure 6C — background noise precluded analysis of fundamental frequency via VocalPy, therefore was estimated manually in Ocenaudio [48]. Spectrograms were generated using the python Matplotlib function `matplotlib.pyplot.specgram()` with the following parameters: NFFT=2048, noverlap=256, Fs=192000.

## 4.5 Data availability

Data and code will be posted publicly upon publication.

## Acknowledgments

REP acknowledges funding support through National Institutes of Health Training Program in Computational Neuroscience 20T90DA059110. ELM received support through the Simons Society of Fellows, as well as the NIH NINDS (award numbers 1K99NS131256-01 and 4R00NS131256-03). AEH would like to acknowledge deutsche forschungsgemeinschaft DFG grant EXC 2117-422037984. We are grateful for funding support to Basis Research Institute from private donations.

## Notes

### Competing Interest Statement

The authors have declared no competing interest.

